# Accelerating CAR T cell manufacturing with an automated next-day process

**DOI:** 10.1101/2024.06.04.596536

**Authors:** Mina Ahmadi, Nicholas Putnam, Max Dotson, Danny Hayoun, Nujhat Fatima, Jasmine Padilla, Gertrude Nonterah, Xavier de Mollerat du Jeu, Yongchang Ji

**Affiliations:** Thermo Fisher Scientific, 5781 Van Allen Way, Carlsbad CA 92008

**Keywords:** CAR T cells, CAR T cell manufacturing, CAR T cell workflow, automation

## Abstract

The traditional method of CAR T cell production involves lengthy *ex vivo* culture times which can result in the reduction of crucial naïve T cell subsets. Moreover, traditional CAR T cell therapy manufacturing processes can prolong time-to-patient and contribute to disease progression. In this study, we describe an innovative and automated 24-hour CAR T manufacturing process that yields a higher percentage of naïve/stem-cell like T cells which have increased cytotoxic activity and cytokine release. The data supports the feasibility of implementing this streamlined manufacturing process in clinics. This approach also has the potential to enhance CAR T therapy efficacy and improve patient access to therapy.

## Introduction

Chimeric antigen receptor-engineered T cells (CAR T) represent a significant breakthrough in the field of cancer treatment.^1,2,3^ While this is a major advancement in treating these otherwise fatal cancers, approximately 30–40% of patients with B cell malignancies show long-term remission upon treatment with CAR T cells highlighting the need for more effective treatments to be developed.^4,5,6^

Furthermore, recent studies have reported that 20-30% of B cell acute lymphoblastic leukemia (B-ALL) patients enrolled for CD19-targeted CAR T cell therapy were unable to receive those treatments due to either disease progression between apheresis and CAR T infusion or manufacturing failures.^7,8,9^

The traditional method of manufacturing CAR T cells involves long *ex vivo* culture times spanning 7 to 14 days which can lead to a reduction in important T cell subsets such as naïve and T memory stem cells. The final product from the traditional method of processing CAR T cells often has an enriched population of highly differentiated T cells. Studies have shown that T memory stem cells (Tscm), which can self-renew and reconstitute different T-cell subsets, are crucial for long-term antitumor efficacy. ^5, 6, 7, 8, 10, 11^ Additionally, some recent preclinical studies demonstrate enhanced effectiveness of CAR T cells against leukemia when they are produced without prior *ex vivo* activation and expansion, or when they are selected from a pool of naive/Tscm precursor cells. Significantly, chronic lymphocytic leukemia (CLL) patients experiencing positive clinical outcomes exhibited a high frequency of naive and Tscm cells in their leukapheresis material. ^12,13,14,15^

Based on preclinical and clinical research highlighting the importance of naive/Tscm cells for sustained antitumor effects, we sought to create a streamlined and automated 24-hour CAR T cell manufacturing process. This innovative approach involves one step isolation and activation of T cells, followed by a 20-hour lentiviral transduction before formulation. By minimizing T cell activation to less than 24 hours, we were able to maintain a significantly higher percentage of naive/stem-cell like T cells, while also accelerating production and cutting costs through automation. Ultimately, our goal is to provide patients quicker access to CAR T cell therapy with enhanced CAR T efficacy. This method, which leveraged FDA CFR part 11 enabled automation software during processing, resulted in highly functional CAR T cells. Our data also supports the feasibility of the manufacturing platform in the clinic.

## Materials and Methods

### CTS™ Cellmation™ Software

CTS Cellmation Software is an automation solution that allows for improved and streamlined control of the cell therapy manufacturing instruments. Cellmation Software works with Thermo Fisher cell therapy instrumentation using batch recipes that are consistent across independent cell therapy workflow runs. The Gibco™ CTS DynaCellect™ system and Gibco CTS Rotea™ Counterflow Centifugation system are equipped with Open Platform Communications United Architecture (OPC-UA) servers which communicate with the instrument-specific modules of the Cellmation Software. Using these modules, CTS Cellmation Software was used to control the T cell isolation process on the CTS DynaCellect system on day 0, as well as the bead removal and wash step on CAR T cell harvest days using the CTS Rotea system as described below (Figure 1).

**Figure 1:**
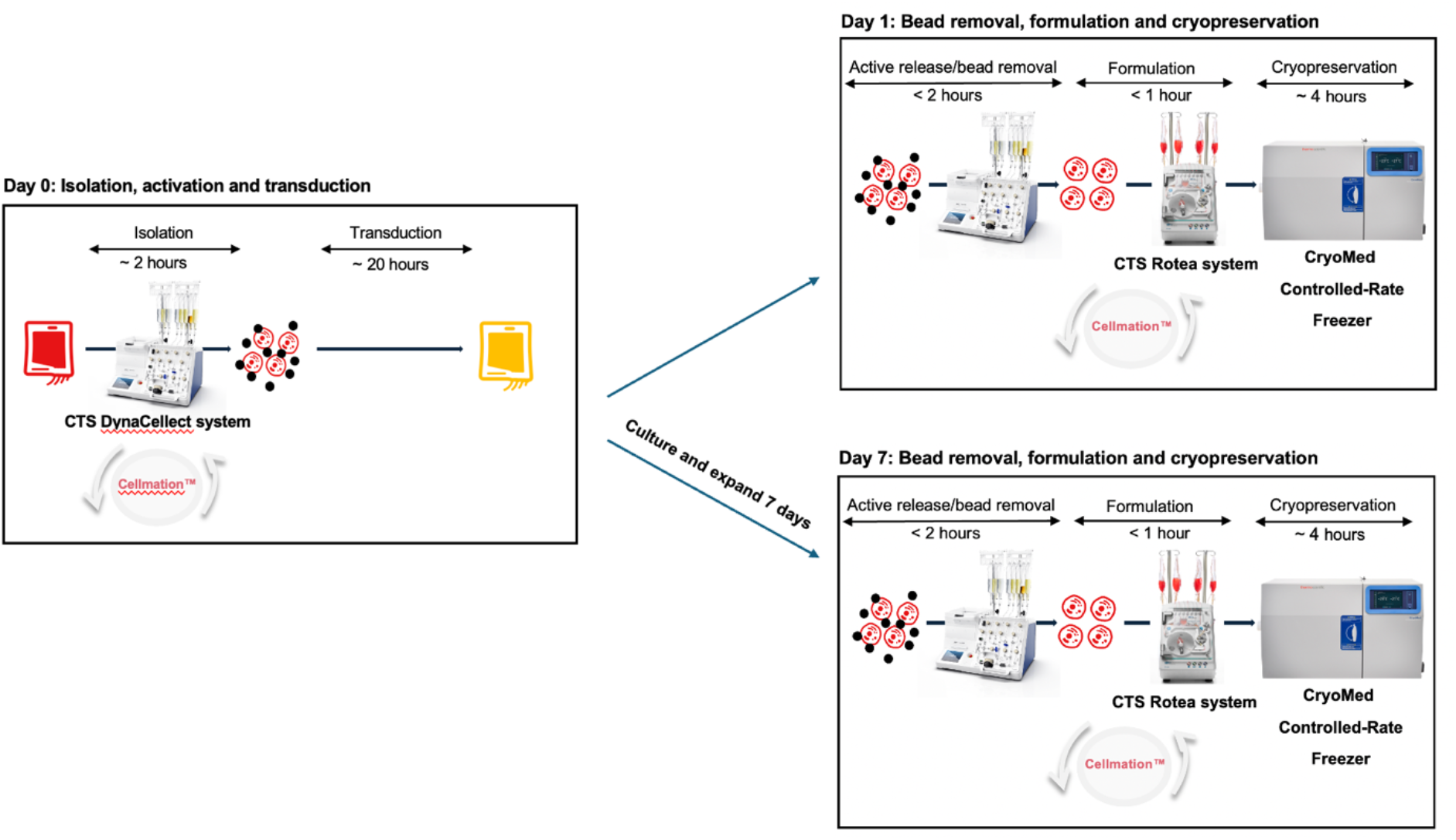
Process flow for 24-hour and 7-day CAR T-cell manufacturing processes. A schematic overview of the shortened 24h workflow and the 7-day workflow after T cells were isolated from the donor’s leukapheresis material, activated, and transduced with a lentiviral vector encoding the CD19-targeting CAR construct. For one batch of cells, after ∼20h culture period, beads were removed; cells were washed, concentrated, formulated, and cryopreserved. For the second batch, ex vivo culture continued for 7 days before bead removal, washing, concentration, formulation and cryopreservation.

### T cell isolation

Briefly, a quarter frozen Leukopak (Allcells) was thawed and transferred to a blood bag. Using the CTS DynaCellect system (Thermo Fisher; cat #A55867) and the CTS Detachable DynaBeads™ CD3/CD28 Kit (Thermo Fisher; cat#A56992) at a 3:1 bead-to-cell ratio, we isolated T cells from the thawed Leukopak (see Table 1 in supplemental methods for set up). Isolated T cells were collected into CTS OpTmizer™ Pro serum free medium (Thermo Fisher; cat# A4966101) supplemented with 2.5% CTS Immune Cell SR (Thermo Fisher; cat# A2596101), 2.3% CTS GlutaMAX-I (Thermo Fisher; cat# 1286001, 1% L-Glutamine (Thermo Fisher; cat# 25030081) and 100U/ml PeproGMP™ Human IL-2 Recombinant Protein (Thermo Fisher; 200-02-1MG). The T cell culture was transferred to G-Rex bioreactor vessels (Wilson Wolf; cat# 80500) at a density of ∼4 × 10^6^ cells/ml.

**Table 1:** CTS DynaCellect system set up for T cell isolation from leukopaks.

| Port | Bag |
| --- | --- |
| A | CTS DynaCollect Isolation Bag |
| B | CTS DPBS/1% HSA |
| C | Output bag |
| D | Not used/empty |
| E | CTS DPBS/1% HSA |
| F | CTS OpTmizer Pro medium with 2.5% ICSR |
| G | Cell input/leukopak cells |
| H | CTS Detachable Dynabeads CD3/CD28 |
| I | Supernatant |

Post isolation, isolation efficiency and culture purity were assessed with the Attune Cytpix flow cytometer (Invitrogen; cat# A48664). Cell counts were performed on the NucleoCounter NC- 200™ System (Chemometec; cat# 991-0203).

### Lentiviral transduction

The anti-CD19 CAR lentivirus (CD19 SCFv-CD3z-41BB) was prepared using the Gibco CTS LV-MAX™ Lentiviral Production System (Thermo Fisher; cat# A35684). The lentivirus was purified, aliquoted and stored at -80°C until ready for use. Prior to use, the anti-CD19 CAR lentivirus was thawed at 4°C. The isolated T cells were then transduced with the lentivirus at an MOI of 2. The T cells were then cultured for 20 hours in a G-Rex bioreactor vessel inside a HeraCell VIOS Incubator (Thermo Fisher; cat# 50145504).

### Post-transduction processing

After 20 hours of incubation, T cells were separated from the magnetic beads with the CTS DynaCellect system using the set up in Table 2 in the supplemental methods.

**Table 2:** CTS DynaCellect Bead Removal set-up on day 1.

| Port | Bag |
| --- | --- |
| A | CTS DynaCollect Isolation Bag |
| B | CTS DPBS/1% HSA |
| C | Output bag |
| D | Not used/empty |
| E | CTS DPBS/1% HSA |
| F | CTS Detachable Dynabeads CD3/C28 Release Buffer |
| G | Input bag (bead-bound cells) |
| H | Not used/empty |
| I | Not used/empty |

Briefly, the transduced T cells were pumped directly from the CTS Dynacellect system into the input bag of CTS Rotea Counterflow Centrifugation System while the magnetic beads remained on the DynaCellect magnet. CAR T cells were then washed using the CTS Rotea™ Counterflow Centrifugation System (Thermo Fisher; cat# A44769) and the CTS Rotea Single Use Kit (Thermo Fisher; cat# A49313) as follows.

We welded an empty 1L bag for waste, a 300mL bag containing 200mL of wash buffer – DPBS (Thermo Fisher; without calcium and magnesium, cat# A1285602) supplemented with 2% HSA (Octapharma; cat# ALB064302) - 1L bag containing the CART T cells and an empty 250mL CryoStore bag (OriGen Biomedical; cat# CS250) to the Rotea Single Use Kit. Table 3 in the supplemental methods shows the ports on the Rotea Single Use Kit each bag was welded onto.

**Table 3:** CTS Rotea system set up for CAR T cell cryopreservation.

| Ports |  |
| --- | --- |
| A | 1000mL bag for waste |
| B | 200mL wash buffer (DPBS/2% HSA) |
| C | Not used/no bag attached |
| D | CAR T cell input loop with G |
| E | Not used/no bag attached |
| F | Not used/no bag attached |
| G | CAR T cell input loop with D |
| H | 250mL CryoStore bag/output |

Using the Rotea system, CAR T cells were washed and then collected and divided into multiple batches for freezing. These batch were collected in 50mL Cryostore bags and frozen in a final 50 × 10^6^ cells/mL cell density for cryopreservation using CS10 Freeze Medium (STEMCELL Technologies; cat#07930). Once CAR T cells had been harvested in CS10 Freeze Medium, the bag was stored in the CTS CryoMed™ Controlled-Rate Freezer (Thermo Fisher; cat# TSCM17PA). Once the storage temperature in the controlled-rate freezer reached -80°C, the bag with CAR T cells was moved into liquid nitrogen for long-term storage.

The cells for 7-day control were cultured (0.5 × 10^6^/cm^2^ cells) for seven days *ex vivo* in CTS OpTmizer Pro medium. This allowed us to compare the differences between the CAR T cells cryopreserved after the 24h manufacturing process versus cells that were further cultured ex vivo for seven days, mimicking a traditional process that includes the expansion step.

### Flow cytometry

Cell samples were collected and centrifuged at 400xg for 5 minutes at room temperature to remove the cell culture or wash buffer medium. The supernatant fraction was removed, and the sample was washed with 1mL eBioscience^TM^ flow cytometry staining buffer (FSS) (Invitrogen; cat# 00-422-26).

The samples were centrifuged again at 400xg for 5 minutes at room temperature followed by removal of the supernatant fraction. The cell sample was then resuspended in a 20% solution of eBioscience Anti-Hu Fc receptor binding inhibitor, (Fc block) (Invitrogen; cat# 14-9161-73) diluted in FSS. Cell samples were plated into 96-well round bottom plates (Costar; cat# 3879). ∼0.3×10^6^ cells were plated per well in a total volume of 100μL. Each sample was divided into 3 groups: stained, unstained plus viability indicator dye, unstained. Incubation in the 20% Fc block solution spanned 15 minutes at 4°C in the dark. After this, the stained groups were incubated in freshly made antibody cocktails for 15 minutes at 4°C in the dark.

Then, the cells were centrifuged again at 400xg for 5 minutes at room temperature. The supernatant fraction was removed, and the samples were washed with 200μL FSS. This was followed by centrifugation at 400xg for 5 minutes at room temperature. The supernatant fraction was removed and the stained and unstained plus viability indictor dye groups were resuspended in 200μL of a 1:500 dilution of the viability dye. Fixable dyes were incubated for 30 minutes at room temperature protected from light. The unstained control groups were resuspended in 200μL of FSS. This was followed by acquisition of the samples on the Attune CytPix flow cytometer with CytKick^TM^ MAX autosampler (Invitrogen; cat# A38976).

Data was acquired with Attune Cytometric Software v6.0.0 and analyzed using FlowLogic 8.6. Results were visualized using GraphPad Prism v9.3.1 (350).

Flow cytometry antibody panels used are outlined in the supplemental methods.

### Cytotoxicity assay

Cytotoxic activity of the CAR T cells was evaluated with the Promega ONE-Glo™Luciferase Assay (Promega Corporation; cat# E6120) according to manufacturer instructions. CAR T cells from the 24-hour process as well as CAR T cells that were cultured ex vivo for an additional 7 days were thawed, cultured for 3 days in CTS OpTmizer Pro serum-free media. The cells were then co-cultured with luciferase-expressing Nalm6 cells for 5 hours. The effector-to-target ratio used in these studies ranged from 10:1 to 0.312:1.

Following incubation and addition of ONE-Glo reagents as described by the manufacturer, luciferase levels were measured using the Varioskan™ microplate reader (Thermo Fisher).

### DNA sample collection

2×10^6^ CAR T cells were collected for DNA extractions. DNA samples were isolated using the PureLink^TM^ Genomic DNA Mini Kit (Thermo Fisher; cat# K182002) under sterile conditions as described by the manufacturer. Each sample was eluted into sterile DNase/RNase-Free 1.5mL microcentrifuge tubes (Thermo Fisher; cat# AM12400) then divided into 50μL aliquots. All samples were stored at -80°C until use.

### RNA sample collection

Cell culture supernatant was collected post-transduction and aliquoted into DNase/RNase-Free 1.5mL microcentrifuge tubes (Invitrogen; cat# AM12400). Samples were stored at -80°C until RNA extraction. RNA was extracted and purified using PrepSEQ Nucleic Acid Sample Preparation kit (Applied Biosystems; cat# A50485) as described by the manufacturer. RNA samples were used within 1 hour of sample purification.

### qPCR assay preparations

#### Mycoplasma identification

Before each assay preparation the biosafety cabinet and pipettes were wiped down with freshly prepared 10% bleach, followed by UV sterilization for 30-60 minutes. The MycoSEQ^TM^ Mycoplasma Real-Time PCR Detection qPCR Kit (Applied Biosystems; cat# 4460623) was used to detect mycoplasma contamination and the assay was performed as described by the manufacturer. Briefly, DNA samples designated for mycoplasma detection were diluted with nuclease free water (Gibco; cat# 10977-015) to obtain 15ng DNA per reaction well. The qPCR premix solution was composed of Power SYBR^TM^ Green PCR Master mix 2x, Mycoplasma Real-Time PCR Primer Mix 10x, nuclease free water, and discriminatory positive control (DPC) or DNA samples. We also included a set of controls without template DNA (NTC) and/or with the DPC.

#### Viral copy number (VCN)

VCN was quantified using the ViralSEQ^TM^ Lentivirus Proviral DNA Titer qPCR Kit (Applied Biosystems; cat# A53562). The assay was performed as described by the manufacturer. 1μL of DNA sample was added to each reaction well. The qPCR premix solution was composed of 2x Environmental Master Mix, Proviral DNA Titer Assay Mix, nuclease free water, and DNA samples. The provided standard was used to create the standard curve used for VCN quantification. Additionally, a set of samples were run without template DNA (NTC).

Once we acquired the data, the equation of the line was generated via a standard curve.

#### Residual lentiviral vector

The ViralSEQ^TM^ Physical Titer Kit (Applied Biosystems; cat# A52597) was used to measure residual lentiviral vectors during CAR T cell generation. Experimental procedure was performed as described by the manufacturer. Briefly, each RT-qPCR reaction contained 5μL of sample RNA. Each reaction was composed of 2X RT-PCR buffer, 25X RT-PCR enzyme mix, physical titer assay mix, nuclease free water, and RNA samples. A provided standard was diluted to produce a standard curve for residual lentiviral particle quantification. We also included a set of samples without template DNA (NTC).

For all qPCR and RT-qPCR assays, premixed solutions were plated in MicroAmp^TM^ Optical 384-Well Reaction Plates with Barcode (Applied Biosystems; cat# 4309849) in triplicates. Plates were sealed with MicroAmp^TM^ Optical Adhesive Film (Applied Biosystems; cat# 4311971) and centrifuged at 1000xg for 2 minutes. All bubbles were removed then the samples were run on the QuantStudio^TM^ 12KFlex (Applied Biosystems; cat# 4470661). Data was acquired with the QuantStudio System.

### Cytokine release

Five hours after the cytotoxicity assay was performed, 180µL of supernatant was collected and pooled from the triplicate in sterile 1.5mL microcentrifuge tubes. Samples were centrifuged at 1,400 rpm for 10 minutes. Samples were stored at -80°C until use.

The ProcartaPlex Hu Th1/Th2/Th9/TH17/TH22/Treg 18plex kit (Invitrogen; cat# EPX180-12165-901) was used to quantify cytokine release. All reagents, standards, and samples were prepared and ran as described by the manufacturer. Samples were analyzed on the Luminex 200 System (Luminex; cat# LX10009295406, LXY09323102) running the Luminex xPONENT for LX100/LX200 software (v.4.3 Update 1). Data was analyzed using the ProcartaPlex web app.

## Results

CAR T cells were produced from healthy donor leukpaks. The T cells were isolated and activated using CTS Detachable CD3/CD28 Dynabeads and the CTS DynaCellect system. Isolation was completed within 2 hours and was followed by transduction of bead-bound cells using a lentiviral vector encoding anti-CD19 CAR at MOI of 2 for approximately 20 hours. To compare CAR T cells produced with the shorter workflow with a conventional CAR T cell manufacturing process that includes an expansion step, after transduction, the cell-bead culture was either harvested after 24 hours or cultured *ex vivo* for 7 days post-transduction (Figure 1).

Immediately after T cell isolation (day 0), we observed an average of 84% isolation efficiency across the three donor leukopaks used in this study (Figure 2A). As shown in Figure 2B, we obtained high T cell recovery on the day of harvest for both the 24-hour and 7-day processes with averages of 82% T cell recovery and 78% T cell recovery for the respective processes. Recovery is a measurement that compares the number of T cells prior to formulation process with the number of T cells recovered post formulation. The cells remained highly viable post-isolation and through the CAR T cell manufacturing process (Figure 2C). We also assessed T cell recovery and viability after thawing both 24-hour and 7-day process products. We observed comparable T cell viability and recovery post-thaw for both products (Supplemental figure 2A-B).

**Figure 2:**
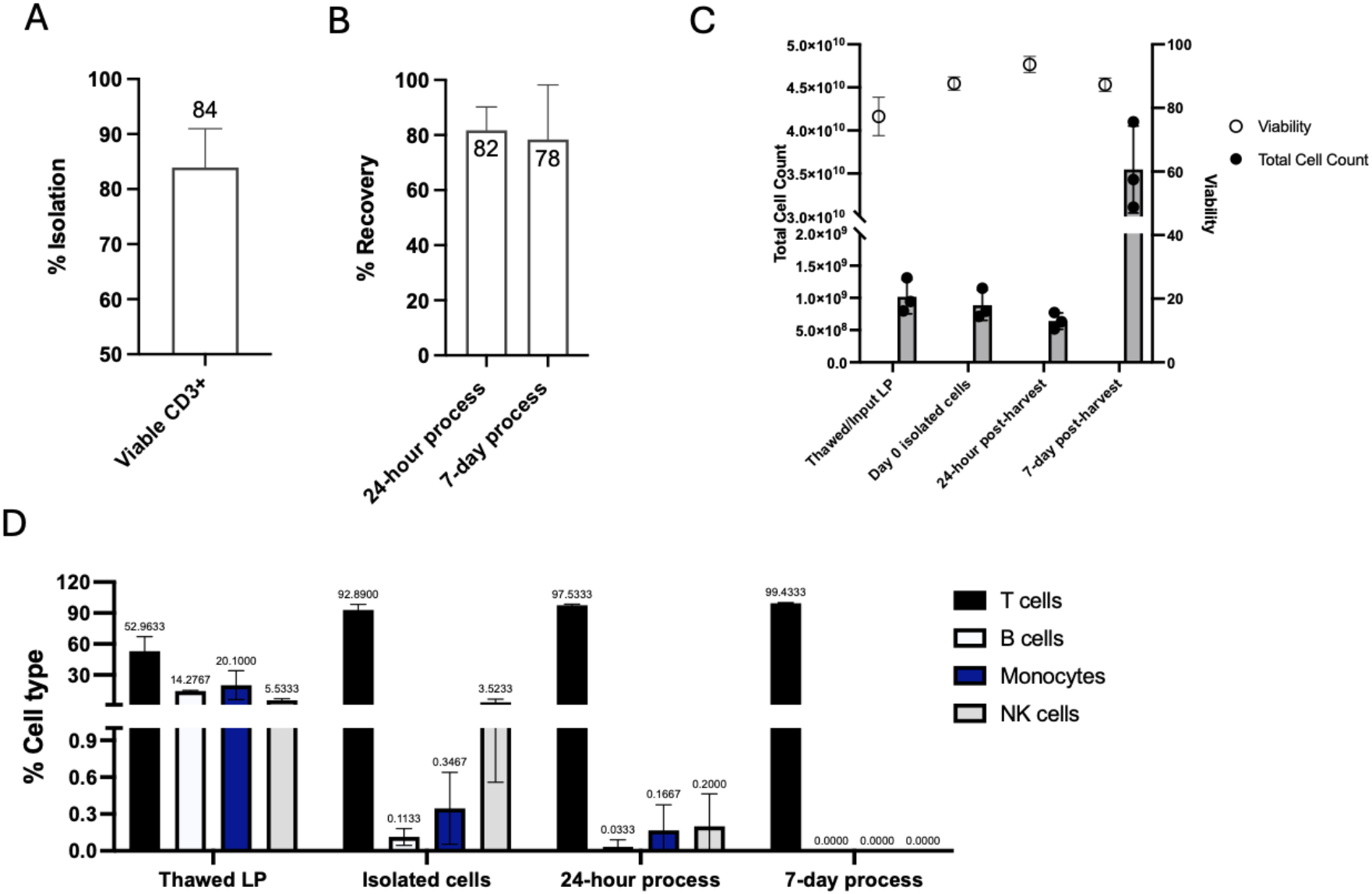
Characterization of T cells on day 0, at 24 hours and 7 days. 2A) T cell isolation efficiency on day 0 for three donors. Isolation efficiency measures how many T cells are in the supernatant versus the T cells in the input sample for isolation. 2B) Comparison of T cell recovery between the 24h and 7-day processes post-harvest (n=3, graph shows mean + SD; unpaired student two-tailed t-test). 2C) Cell viability and total T cell count on leukapheresis material, day 0 isolated T cell culture, harvested 24h and 7-day CAR T cells (n=3, graph shows mean + SD). 2D) Immuophenotypes of the leukapheresis material and isolated T cell culture on day 0, as well as the 24h time-point and the day 7 time point prior to cryopreservation, respectively (n=3, graph shows mean + SD; unpaired student two-tailed t-test).

Cell viability remained high (>85%) throughout the CAR T cell manufacturing process (Figure 2C). We observed ∼55-fold increase in total T cell content from 24-hour to 7-day process due to extended ex vivo culture (Figure 2C).

Flow cytometry was used to assess immune cell composition throughout the process. Our day 0 isolated culture as well as the final 24-hour and 7-day products showed greater than 95% T cell purity with a distinct increase in T cell content post-day-0-isolation (Figure 2D).

NKT (CD3^+^CD56^+^) cells were the major non-T cell fraction isolated using this process. This is anticipated given that NKT cells express CD3 which enables their isolation using detachable CD3/CD28 beads (Supplemental Figure 1A). However, we did observe a significant decrease of CD56+ cells (NK and NKT) between day 0 isolated culture and 24 hours later when CAR T cells were harvested (Figure 2D).

We assessed the expression of early (CD69), mid (CD25), and late activation marker (HLA-DR) on days 0, 1, 2 and 3 using flow cytometry. CD69 showed peak expression on day 1, while CD25 peaked on day 2, and HLA-DR on day 3 as anticipated (Supplemental Figure 1B).

Overall, this data demonstrates that our T cell isolation process was highly efficient resulting in viable isolated cells that could be used for various downstream applications.

The CAR expression levels were determined prior to harvesting and formulation through immunostaining using a monoclonal antibody that specifically targets the idiotype of the single-chain variable fragment. We observed an average of 76.4% CAR expression in the product generated through 24-hour process whereas that number dropped significantly to 12.2% when these CAR expressing cells were cultured *ex vivo* for an additional 7 days (Figures 3A and 3B). This observation is likely due to pseudo-transduction that occurs post-transduction, given the naiver state of the T cells at the time of transduction as reported previously.^12^ Membrane proteins expressed in packaging cells can passively be transferred to lentivirus envelope during LV production.^16,17^ These proteins can then be passively transferred to the T cell membrane post transduction. While this phenomenon happens in both naive and active T cells, higher rates of cell membrane turnover in activated T cells could lead to more efficient the removal of membrane proteins that were passively transferred and potentially contributing to pseudotransduction.^18^ Due to the brief culture time, the 24-hour CAR T cells did not exhibit robust expansion. In comparison, there was a 52.6-fold average fold expansion for CAR T cells manufactured using the 7-day process (Figure 3D). This resulted in an average of 520 million and 3.4 billion CAR-expressing T cells for 24-hour and 7-day process, respectively (Figure 3C). Despite a decrease in the percentage of CAR-expressing cells due to the removal of pseudo-transduced cells during the 7-day ex-vivo culture process, the high cell expansion allowed for the generation of 3.4 billion CAR T cells on day 7.

**Figure 3:**
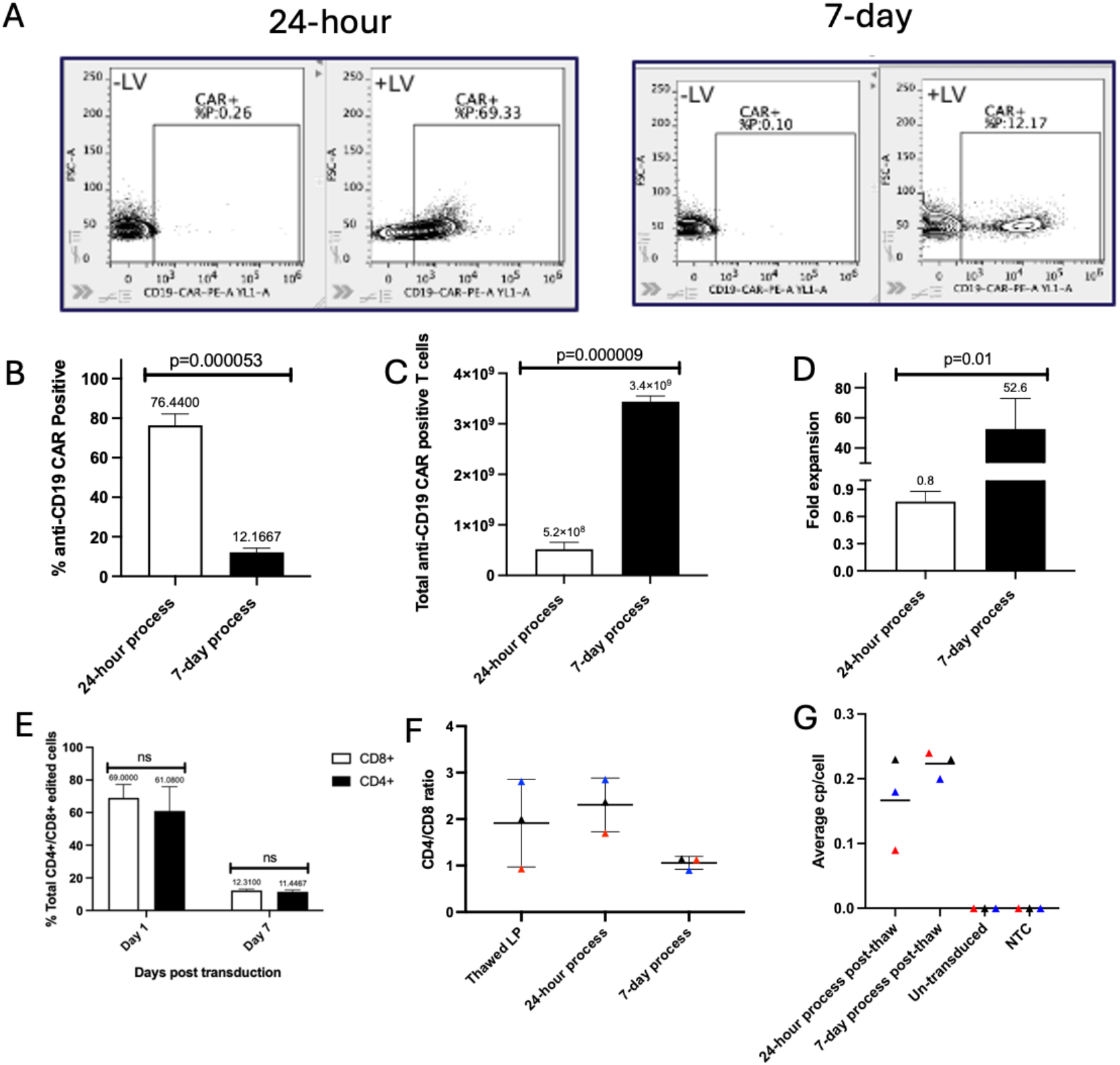
Characterization of products generated through 24-hour and 7-day processes prior to cryopreservation. 3A) and 3B) demonstrate a comparison of percent CAR expression on products from 24-hour and 7-day processes as measured by flow cytometry (n=3, graph shows mean + SD; unpaired student two-tailed t-test). 3C) and 3D) show a comparison of T cell expansion between the 24-hour and 7-day processes (n=3, graph shows mean + SD; unpaired student two-tailed t-test). While 3E) shows a comparison of transduction efficiency in CD4 and CD8 T cells on day 1 and day 7 prior to harvest for the 24-hour and 7-day processes, respectively (n=3, graph shows mean + SD; two-way ANOVA with Sidak’s multiple comparisons test). 3F) shows the CD4:CD8 ratio on day 0, day 1 and day 7 (n = 3; Graph shows individual data points with mean + SD). 3G) The average vector copy number (VCN) post-transduction was generated using the ViralSEQ^TM^ Lentivirus Proviral DNA Titer qPCR Kit. Results were set relative to the number of cells in each reaction well. The 24-hour and 7-day process resulted average cps/cell > 0.1 copies (24-hour process = 0.17 average cps/cell, 7-day process = 0.22 average cps/cell) with no significant difference between the means. *P = 0.2550*. The un-transduced and NTC controls showed no cps/cell.

We then analyzed transduction efficiency in CD4 and CD8 T cell types. While there were no significant differences in transduction between CD8+ and CD4+ T cells, transduction was only slightly higher in CD8+ T cells on the day of harvest for both processes (Figure 3E). The CD4:CD8 ratio was also determined by analyzing the percentage of CD4+ and CD8+ T cells. We calculated the CD4:CD8 ratio based on the percentage of CD4+ and CD8+ T cells present in the thawed leukopak, at 24 hours, and for the 7-day process. While the ratio averaged at approximately 2 for the 24-hour CAR T cells, the ratio came down to 1 at the end of the 7-day process (Figure 3F). An analysis of lentiviral copy number revealed that the 24-hour and the 7-day CAR T processes yielded similar levels of CD19 CAR gene copies, as measured by qPCR. This observation further establishes the point that transduction of T cells using the shorter 24-hour workflow is efficient and can be used in manufacturing functional CAR T cell therapies (Figure 3G).

To demonstrate the pharmacologic activity of anti-CD19 CAR T-cells generated using 24-hour vs 7-day processes, we performed phenotypic and functional characterization experiments. To do so, we first thawed the end products from both processes and cultured the CAR T cells for 72 hours.

Firstly, we investigated the *ex vivo* proliferation of 24-hour and 7-day CAR T cells by thawing frozen CAR T products from each process and stimulating them with 200U/ml IL2. We then tracked cell expansion during culture. We observed that the 24-hour CAR T cells proliferated faster compared to the 7-day CAR T cells. By day 4, the 24-hour CAR T cells expanded by ∼9.50 fold, while cell generated through 7-day process only expanded by ∼5.25 fold (Supplemental Figure 2C).

We also characterized the memory phenotype composition of final products of 24-hour and 7-day process 3 days post thaw using flow cytometry. Consistent with previous reports,^10,11,12^ we observed that compared to standard 7-day process, the 24-hour CAR T cell product showed a less mature T-cell memory profile that closely resembled the phenotype of the starting material. This suggests that shortening ex vivo culture time leads to a product with less differentiated T cells as shown in Figures 4A-4C. Furthermore, we also detected a higher percentage of PD-1+ and LAG3+ T cells for the 7-day process product compared to 24-hour product (Figure 4D). Together these data indicate that the 24-hour CAR T cells exhibited a more naïve T cell phenotype and were less exhausted compared to the 7-day CAR T cells.

**Figure 4:**
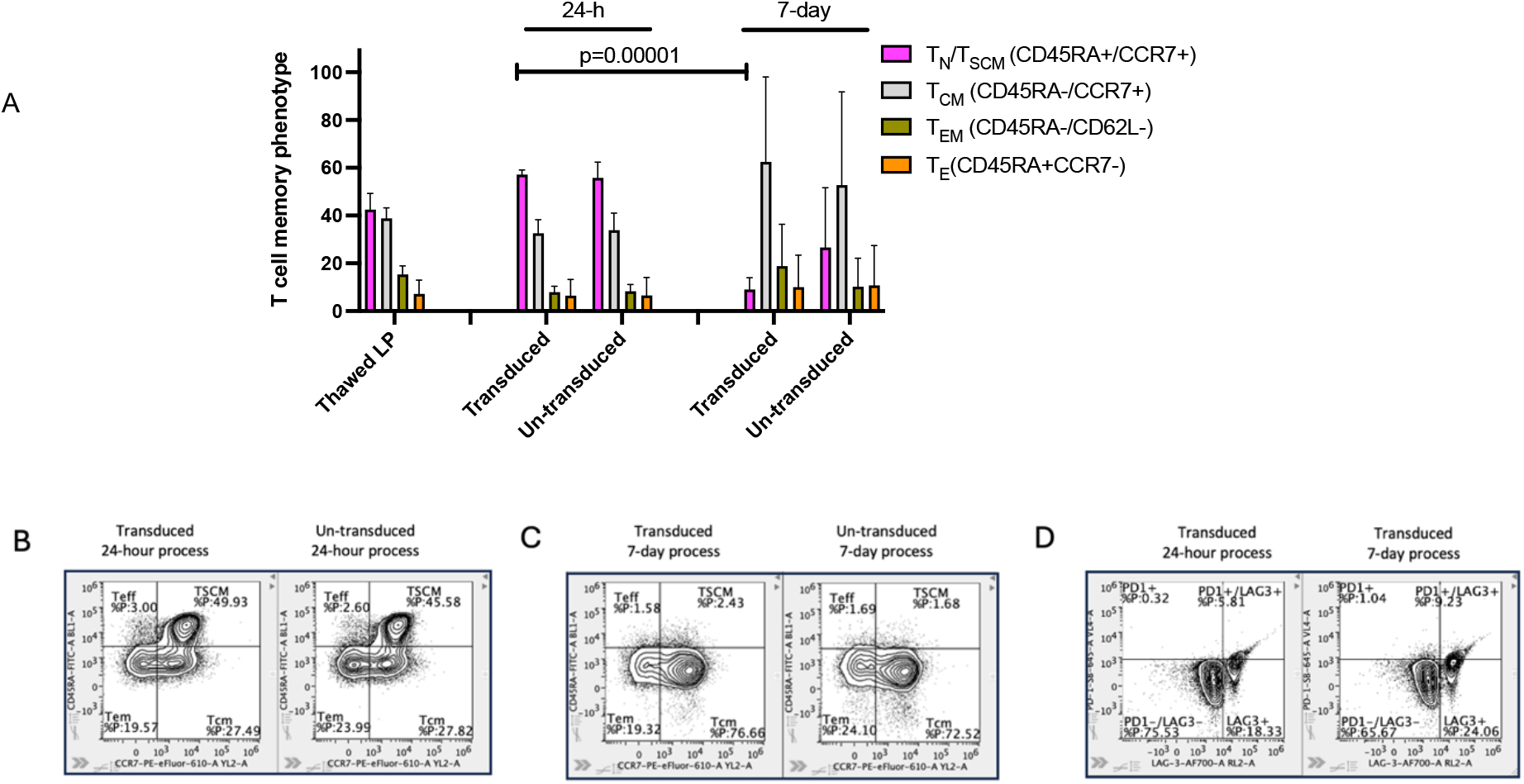
Characterization of thawed products generated through 24-hour and 7-day processes prior to cryopreservation. 4A-C) demonstrate a comparison of T cell memory phenotypes from products generated using 24-hour and 7-day processes by flow cytometry, 3 days post-thaw (n=3, graph shows mean + SD). 4D) looks at the T cell exhaustion marker (PD1 and LAG3) expression between the 24-hour CAR T cells and the 7-day CAR T cells 3 days post-thaw (n=3, graph shows mean + SD).

We then investigated if maintaining the less differentiated state of CAR T cells impacts their functionality. To do this, we performed target-specific cytotoxicity against CD19-expressing NALM6 cells. The cytotoxic activity of the 24-hour CAR T cells trended higher than the 7-day CAR T cells indicating potentially higher potency (at lower effector: target ratios), most likely due to their less differentiated memory phenotype (Figure 5A).

**Figure 5:**
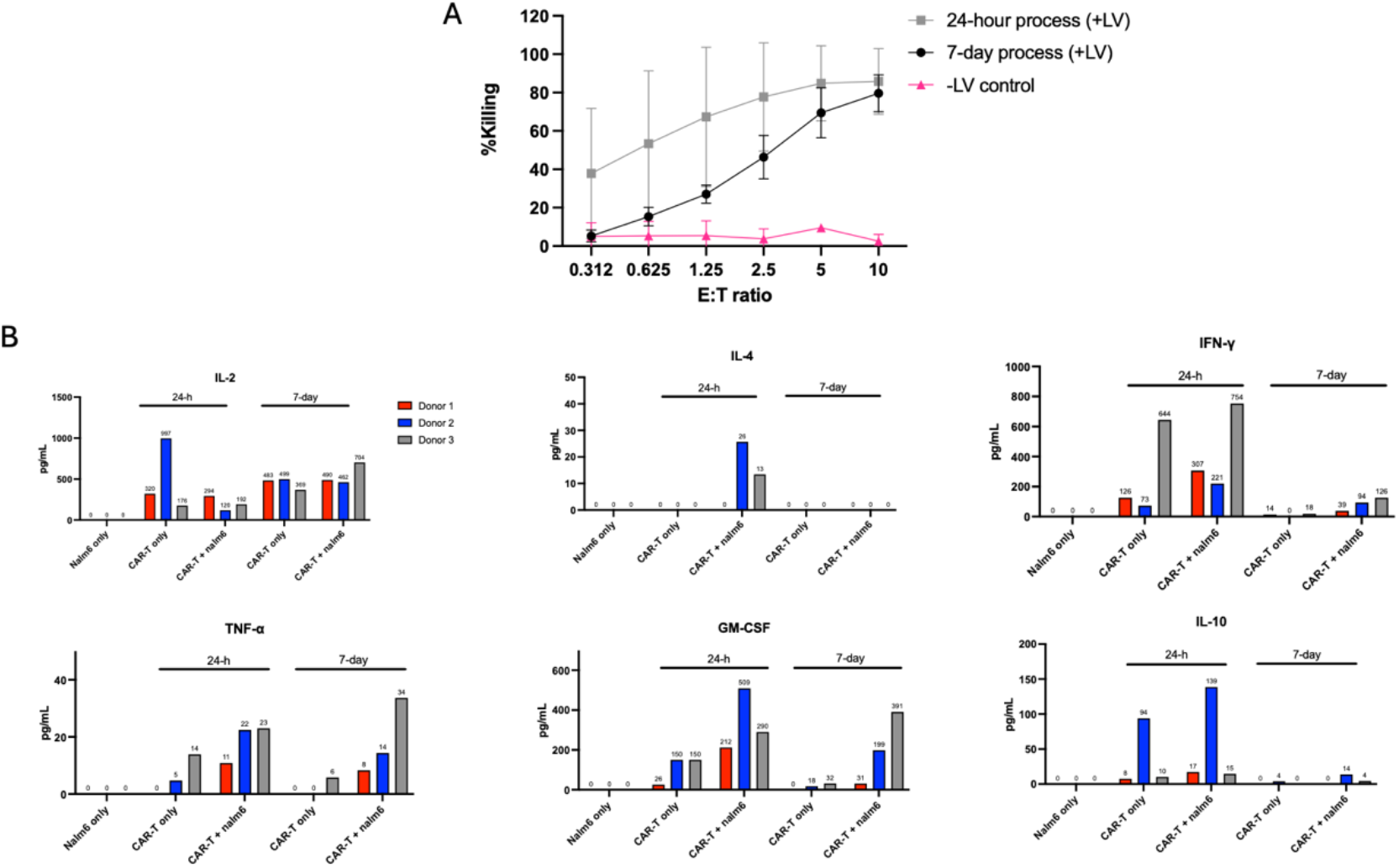
Cytotoxic killing and quantification of cytokines release post cytotoxic killing. 5A) showcases cytotoxicity of 24-hour and 7-day CAR T cells exposed to CD19+ target Nalm6-Luc cells for 5 hours. Target cell killing efficacy was calculated by measuring luciferase activity. (n=3, graph shows mean + SD, unpaired two-tailed student t-test). For 5B) we used the ProcartaPlex Hu Th1/Th2/Th9/TH17/TH22/Treg 18plex Kit to measure cytokine release. IL-4, IL-10, IFN-γ, IL-1β, GM-CSF, and TNF-α concentrations post cytotoxic killing showed no significant difference between 24-hour and 7-day process (IL-4, *P=0.1537*; IL-10, *P=0.2844*; IFN-γ, *P=0.1109*; IL-1β, *P=N/A*; GM-CSF, *0.3940*; TNF-α, *P=0.9988*). A significant difference between the 24-hour process and 7-day process was observed with IL-2 (*P=0.0188*) release post cytotoxic killing.

To better characterize potency, we studied cytokine release from the CAR T cells 5 hours after they were incubated with CD19-expressing Nalm6 cells. Generally, while the difference in cytokine release between 24-hour CAR T and 7-day CAR T cells was not statistically significant, 24-hour CAR T cells tended to secrete higher levels of IL-4, IL-10, GMCSF and IFN-γ (Figure 5B). Our samples that were used for cytokine were collected from wells containing a 10:1 effector-to-target ratio and the results of the cytokine assay were consistent with the higher trend in cytotoxicity activity we observed with the 24-hour CAR T cells.

We tested our final product for mycoplasma and residual virus using PCR-based assays as instructed by the manufacturer. We measured mycoplasma levels in the final CAR T products using qPCR. CT values were greater than 35 suggesting no mycoplasma contamination in the final CAR T product (Supplemental Figure 3A). We also measured residual lentiviral particles pre- and post-formulation on day 1 and day 7 for 24-hour and 7-day processes via qPCR. Donors 2 and 3 showed large reductions in residual viral particles post-formulation during the 24-hour process (Donor 2 = 88-fold reduction, Donor 3 = 10-fold). The pre-formulation sample for Donor 1 was not collected. However, the post-formulation value for Donor 1 fell below the values observed for Donors 2 and 3. The 7-day process showed a similar reduction trend of the residual viral particles across the donors (Donor 2 = 50-fold reduction, Donor 3= 2.9-fold reduction). As observed with the 24-hour process, Donor 1 showed reduced residual vector relative to the other donors (Supplemental Figure 3B)

## Discussion

The traditional autologous CAR T cell therapy process involves several steps: collecting T cells from the patient, genetically modifying them to express CARs, expanding the modified T cells in culture, and then harvesting, formulating, and cryopreserving them. This is followed by characterization and re-infusion into the patient to target and eliminate cancer cells. The entire process typically takes around 3-4 weeks, including preparation time. However, the production of these CAR T cell therapies is a costly endeavor. Furthermore, the lengthy manufacturing process can lead to a reduction in the potency of the CAR T cells due to their terminal differentiation and exhaustion. As a result, while CD19-targeted CAR T cell therapy is highly effective in treating B cell malignancies, these challenges can limit its widespread adoption.

In this study we highlighted an innovative 24-hour process that streamlines the production of vein-to-vein CAR T cell therapies generated using an anti-CD19 encoding lentivirus. This shortened manufacturing process improves patient’s access to therapy, reduces expensive manufacturing time, and enhances the potency of CAR T cells, resulting in a high-quality product with demonstrated target-specific cytotoxicity. By leveraging rapid manufacturing process, our process outpaces traditional methods, which are often labor-intensive and time-consuming.

Furthermore, this process utilizes an array of automated and closed instruments which are digitally integrated leading to minimal human touch points. Being able to automate this process is crucial as automation can reduce the risk for contamination and error that can arise from manual handling of cells and equipment during the CAR T cell manufacturing process.

The 24-hour manufacturing process showed comparable or superior cell viability, T cell recovery post formulation, and T cell purity throughout the process to anti-CD19 CAR T cells derived from conventional manufacturing process.

Although CAR T cells transduced using the 24-hour process demonstrated higher transduction efficiency than those produced using the 7-day process, we believe that this difference is likely due to pseudo-transduction, which is a phenomenon where viral particles bind to cells without actual gene transfer. This suspicion is supported by the observation that the transduction efficiency decreased over time when these cells were cultured, whereas the growth rate of transduced and untransduced cells remained similar. Pseudo-transduction has been shown to be more pronounced in non-activated or minimally activated T cells, although it has been observed in activated T cells as well.^12^ Since cells are simultaneously activated and transduced, most of the cells for the 24-hour process are still “naïve” during transduction process, and this might have encouraged increased levels of pseudo-transduction. Nonetheless, the total numbers of CAR T cells were high for both processes. Understandably, the 7-day process allowed for a greater than 50-fold expansion in CAR T cells. This is probably the pattern that will be observed when 24-hour CAR T cells are infused into patients: a high pseudotransduction rate which tapers off while T cells that are fully/stably edited expand upon exposure to antigen-presenting cells. One significant obstacle that arises due to pseudotransduction is determining the optimal dose. As a result, conventional methods of assessing CAR protein expression, such as flow cytometry-based surface expression, may not be sufficient to accurately evaluate the quality of the transduction process. To better assess the quality of the transduction process, more quantitative measures like Alu repeat-based qPCR may be a more effective approach. However, further research is required to thoroughly validate this method and its effectiveness.

24-hour CAR T cells had increased levels of stem cell-like memory cells (Tscm) over CAR T cells produced using the 7-day process. And likely, due to this “younger” CAR T cell phenotype, there was reduced expression of exhaustion markers^19^ which likely contributed to more efficient killing and cytokine release. This characteristic may have clinical implications, as less differentiated T cells are associated with enhanced *in vivo* expansion and efficacy.

Overall, the data suggests that 24-hour CAR T cells can be manufactured using the process described. Furthermore, 24-hour CAR T cells – which comprised of less exhausted and less differentiated T cells – exhibited equal or superior cytotoxic activity in various effector-to-target-cell ratios.

In conclusion, producing highly functional and more cost-effective CAR T cells has significant implications for the treatment of diseases where CAR T cell therapy has shown to be effective. Minimizing ex vivo culture in combination with automating and closing the manufacturing process, not only enables faster access of patients to these therapies but also enables reducing costs and the rate of process failure. Furthermore, simplified processes have the increased potential to decentralize CAR T cell manufacturing to local hospital laboratories.

## Supplemental Figures

**Supplemental figure 1.**
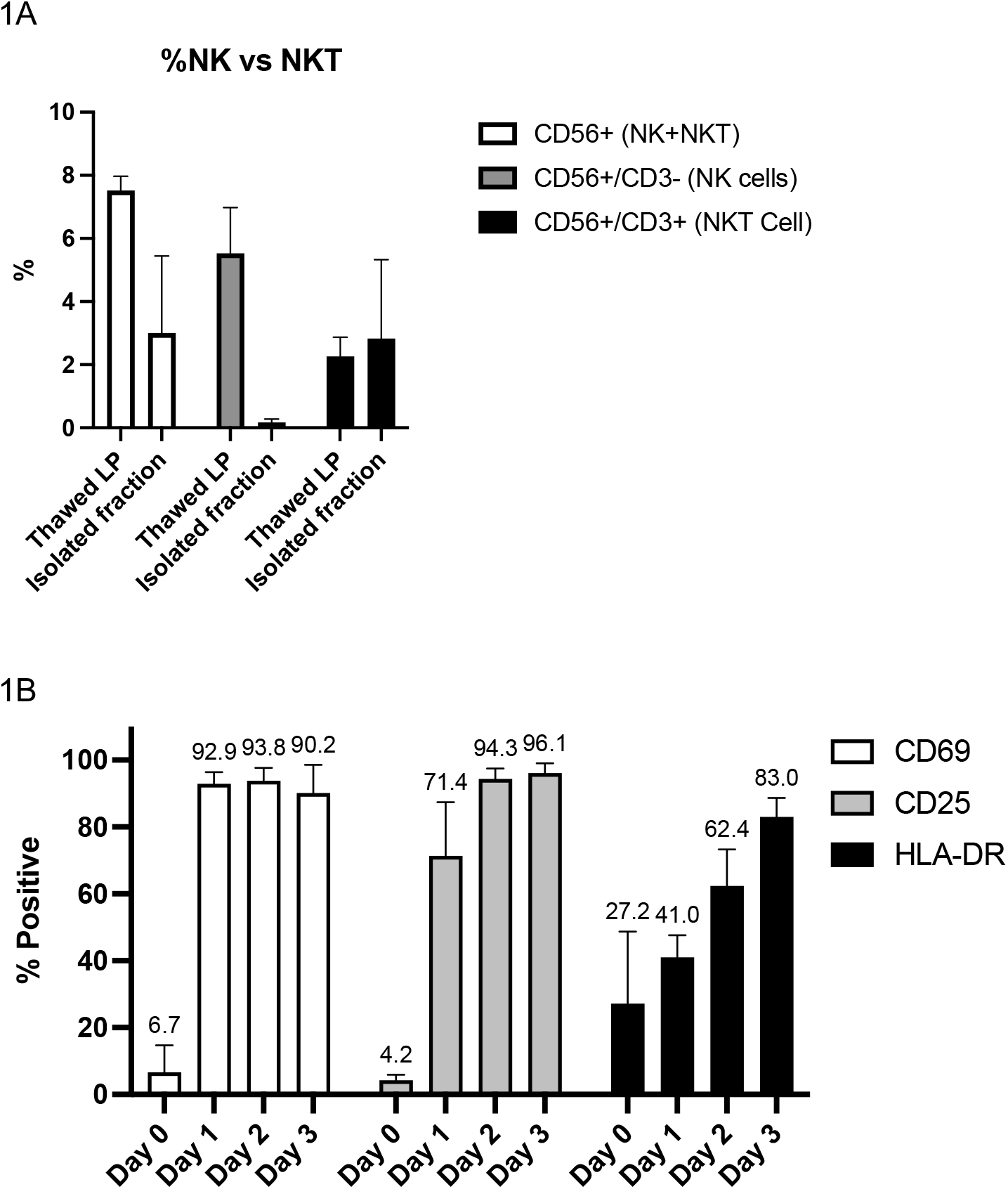
Characterizing activation status of T cells post isolation. Supplemental figure 1. Characterizing isolated T Cells. 1A) Comparing % CD56+ (NK+NKT), vs CD56+/CD3-(NK cells) vs CD56+/CD3+ (NKT cells) in starting leukapheresis material and isolated T cell cultures on day 0. 1B) Expression of early (CD69), mid (CD25) and late (HLA-DR) activation markers post isolation in leukapheresis material and isolated T cell using flow cytometry (n=3, graph shows mean + SD).

**Supplemental figure 2.**
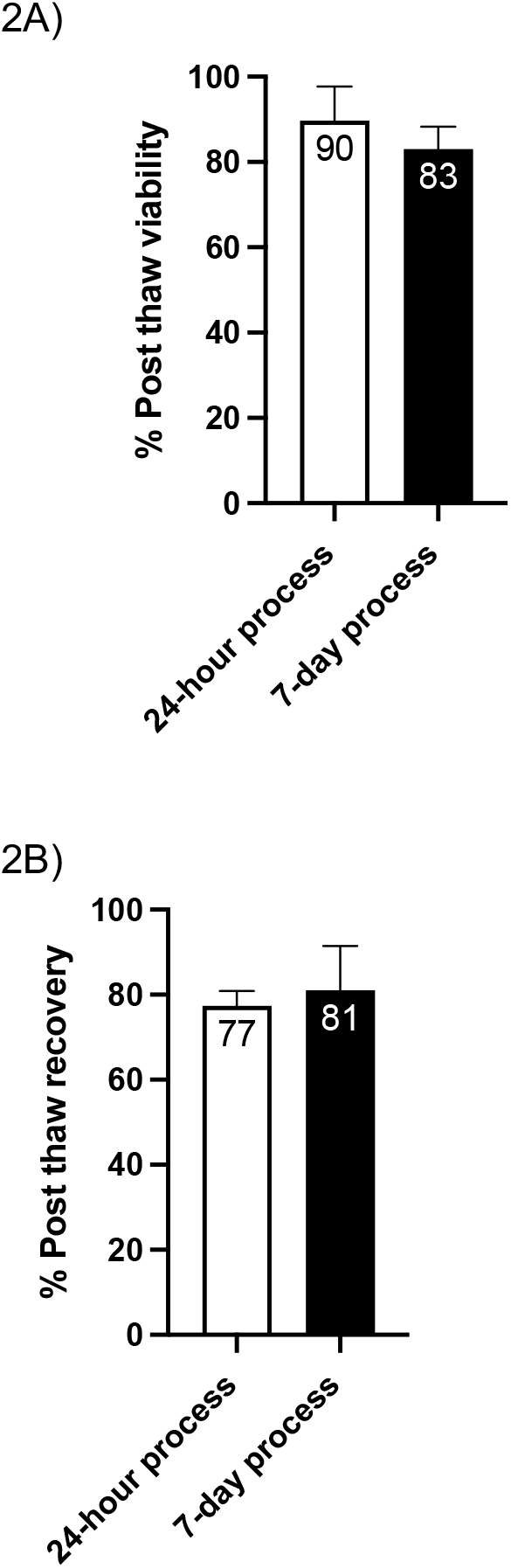
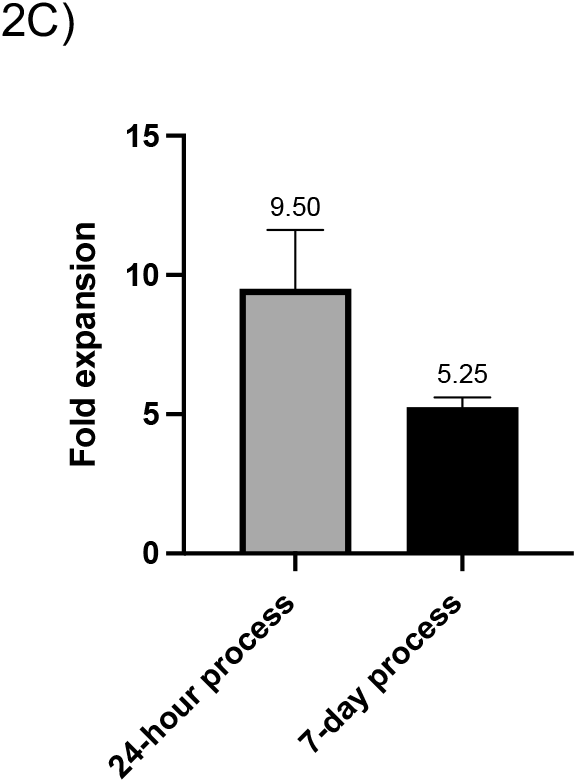
Characterizing 24-hour and 7-day process post-thaw recovery. 2A) and 2B) Comparison of viability and T cell recovery post thaw in 24-hour and 7-day products post-thaw (n=3, graph shows mean + SD). 2C) Comparison of T cell growth 4 days post thaw in 24-hour and 7-day products

**Supplemental Figure 3:**
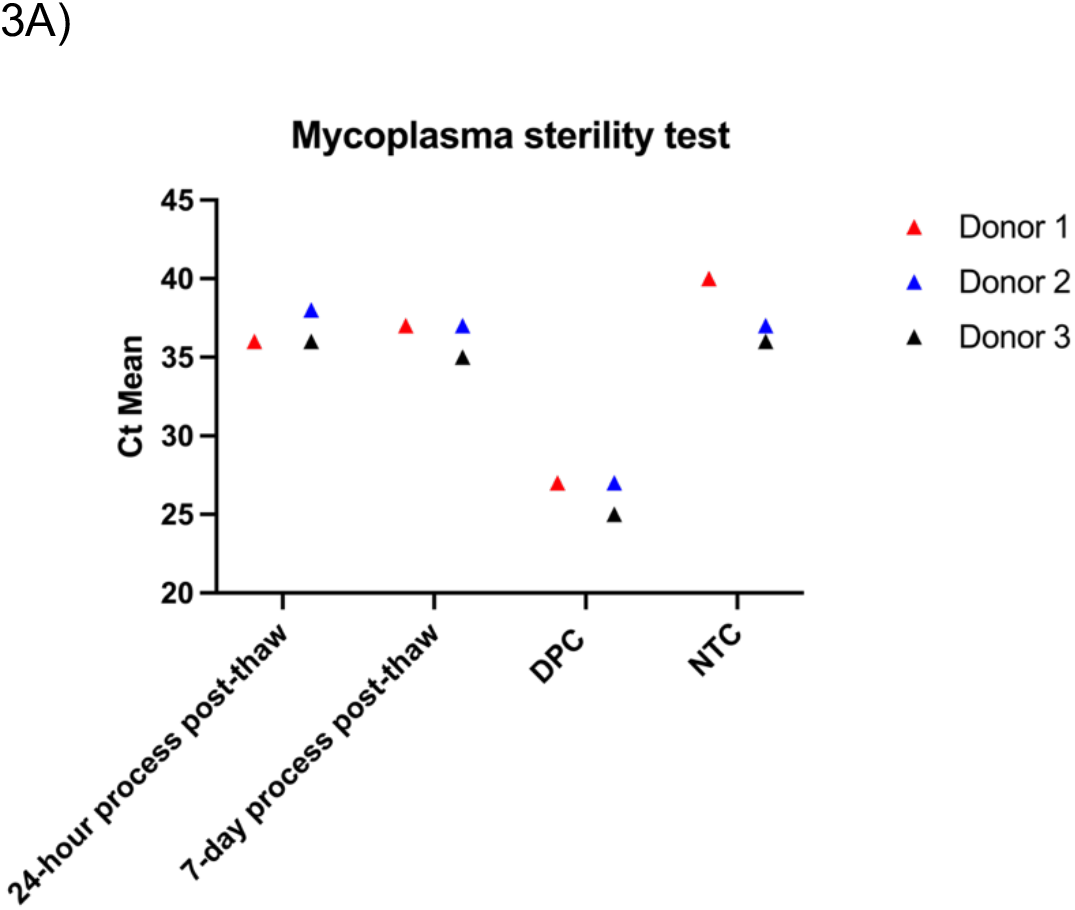
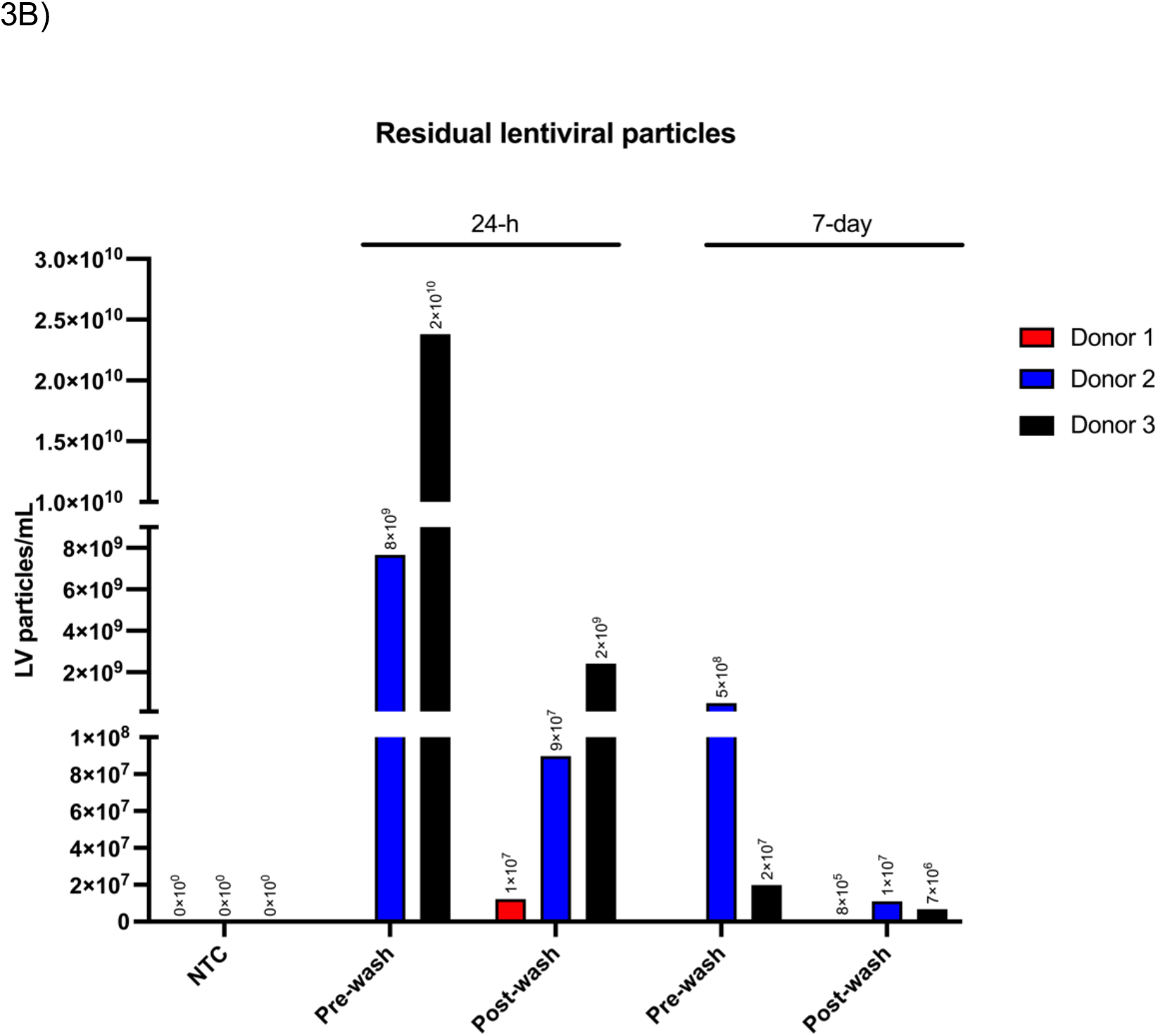
(A) Qualification of mycoplasma free final CAR T product was established using the MycoSEQTM Mycoplasma Real-Time PCR Detection qPCR Kit. Samples were collected three days post-thaw. Ct mean values were used to determine whether contamination was present in the sample. Ct mean < 35 were designated at contaminates. All samples resulted in Ct mean values > 35, suggesting no mycoplasma contamination was present in the final CAR T product. (B) Residual lentiviral particles pre- and post-wash were quantified using the ViralSEQ^TM^ Lentivirus Physical Titer RT-qPCR kit. Donor 2 and 3 showed large reductions in residual viral particles post-washing during the 24-hour process (Donor 2 = 88-fold reduction, Donor 3 = 10-fold). Donor 1 pre-wash sample was not collected; however, the post-wash value falls below with what was observed between Donors 2 and 3. The 7-day process showed a similar reduction trend of the residual viral particles across the donors (Donor 2 = 50-fold reduction, Donor 3= 2.9-fold reduction). Like observed with the 24-hour process Donor 1 showed reduced residual vector relative to the other donors.

## Supplemental methods

### Antibody panels

#### Phenotype

CD3 (Invitrogen; cat# 47-0036-42), CD4 (Invitrogen; cat# 67-0049-42), CD8 (Invitrogen; cat# 56-0087-42), CD14 (Invitrogen; cat# 25-0149-42), CD45 (Invitrogen; cat# 63-9459-42), CD56 (Invitrogen; cat# 12-0567-42), CD19 (Invitrogen; cat# 46-0198-42), CD28 (Invitrogen; cat# 17-0289-42), CD2 (Invitrogen; cat# 78-0029-42), SYTOX^TM^ Blue dead cell stain (Invitrogen; cat# S34857)

#### Activation

CD3 (Invitrogen; cat# 47-0036-42), CD4 (Invitrogen; cat# 67-0049-42), CD8 (Invitrogen; cat# 56-0087-42), CD25 (Invitrogen; cat# 61-0257-42), CD69 (Invitrogen; cat# 46-0699-42), HLA-DR (Invitrogen; cat# 63-9956-42), CD2 (Invitrogen; cat# 78-0029-42), CD56 (Invitrogen; cat# 12-0567-42), SYTOX^TM^ Blue dead cell stain (Invitrogen; cat# S34857)

#### Memory phenotypes

CD45RA (Invitrogen; cat# 11-9979-42), CD62L (Invitrogen; cat# 17-0629-42), CD8 (Invitrogen; cat# A15448), TCR α/β (Invitrogen; cat# 48-9986-42), CD197 (CCR7) (Invitrogen; cat# 61-1979-42), CD4 (Invitrogen; cat# 25-0049-42), V5-Tag (Invitrogen; cat# 12-6796-42), LIVE/DEAD^TM^ fixable aqua dead cell stain kit (Invitrogen; cat# L34966)

#### Exhaustion

CD223 (LAG-3) (Invitrogen; cat# 56-2239-42), CD279 (PD-1) (Invitrogen; cat# 64-9969-42), CD4 (Invitrogen; cat# 25-0049-42), CD8 (Invitrogen; cat# A15448), LIVE/DEAD^TM^ fixable aqua dead cell stain kit (Invitrogen; cat# L34966)

## Author contributions

- MA, XdeM and YJ developed the concept and designed experiments.
- MA, NP, MD, DH, NF, and JP performed the experiments.
- GN prepared the final manuscript.

